# The draft genome of an octocoral, *Dendronephthya gigantea*

**DOI:** 10.1101/487058

**Authors:** Yeonsu Jeon, Seung Gu Park, Nayun Lee, Jessica A. Weber, Hui-Su Kim, Sung-Jin Hwang, Seonock Woo, Hak-min Kim, Youngjune Bhak, Sungwon Jeon, Nayoung Lee, Yejin Jo, Asta Blazyte, Taewoo Ryu, Yun Sung Cho, Hyunho Kim, Jung-Hyun Lee, Hyung-Soon Yim, Jong Bhak, Seungshic Yum

## Abstract

**Background:** Coral reefs composed of stony corals are threatened by global marine environmental changes. However, soft coral communities composed of octocorallian species, appear more resilient. The genomes of several species of cnidarians have been published, including stony corals, sea anemones, and hydra, but as of yet no octocoral species. To fill this phylogenetic gap within the cnidarian, we sequenced the octocoral, *Dendronephthya gigantea,* a non-symbiotic soft coral, commonly known as the carnation coral.

**Findings:** The *D. gigantea* genome size is approximately 276 Mb. A high-quality genome assembly was constructed using 29.85Gb (108x coverage) of PacBio long reads and 35.54Gb (128x coverage) of Illumina short paired-end reads resulting in the largest N50 value reported among cnidarian of 1.4 Mb. About 12 % of the genome consisted of repetitive elements. We found 28,879 protein-coding genes. This gene set contained about 94% metazoan single-copy orthologs, indicating the gene models were predicted with high quality compared to other cnidarians. Based on molecular phylogenetic analysis, octocoral and hexacoral divergence occurred approximately 544 million years ago. Moreover, there is a clear difference in Hox gene composition: unlike in hexacorals, Antp superclass member Evx gene was absent in *D. gigantea.*

**Conclusions:** We present the first genome assembly of a non-symbiotic octocoral, *D. gigantea* to aid in the comparative genomic analysis of cnidarians, including comparisons of stony and soft corals and symbiotic and non-symbiotic corals. In addition, the genome of this species may provide clues about differential genetic coping mechanisms between soft and stony coral regarding the global warming.

## Data Description

### Introduction

Corals, Anthozoa of the phylum Cnidaria, provide habitats for a diversity of marine organisms [1] and are foundational members of the benthic community playing a major role in energy transfer between plankton and the benthos [2]. Corals capture large quantities of plankton and thereby regulate the primary and secondary production of the coastal food chains [2, 3]. Corals can be classified into hexacorals (stony corals and sea anemones) and octocorals (soft corals and sea fans). Global marine environmental changes, represented by the seawater temperature rise and ocean acidification, are known to threaten coral reefs consisting of stony corals in tropical regions [4, 5]. However, soft coral communities in temperate and subtropical regions, seem to prosper owing to their ability to disperse north as distribution limits extend [6, 7]. To date much research has been carried out on the stony corals because of their susceptibility to coral bleaching [5] due to global warming and ocean acidification [8–11]. However, soft corals, which have sclerites, are less vulnerable to such environmental changes [8] and it is suggested that temperate shallow-living octocorals are able to withstand increased levels of temperature and acidification [12]. Although there are significant biological differences between the stony and soft corals in terms of calcification and survival strategies in the changing environment, only hexacoral genomes have been sequenced and analyzed [13–18].

Here, we report the first genome assembly of an octocoral, *Dendronephthya gigantea,* commonly known as carnation coral. *D. gigantea* is a dominant species in the most southern coastal part of Korea [19], in temperate and subtropical regions where yearly water temperature ranges from 14 to 26°C [19]. In general, colonies of this species inhabit shallow water from 10 to 20 m in depth. It is an independent non-symbiotic gonochoric internal brooder. It preys on zooplankton and phytoplankton and does not possess zooxanthellae [20]. These characteristics contrast to those of reef-building *Acropora* species. Our draft genome may therefore serve as a reference for evolutionary studies of azooxanthellate octocorals in terms of understanding different coping mechanisms mediating against rapid environmental changes in comparison to published genomes of stony corals.

### Sample collection and DNA / RNA extraction

A *D. gigantea* colony was collected at approximately 20 m underwater near Seogwipo, Jeju Island, South Korea (33° 13'39'' N, 126° 34'03’' E) on May 22, 2015 using standard scuba techniques (Figure 1A). The underwater yearly temperature range of the site was measured to be 15 and 26 °C. The colony of *D. gigantea,* which carry mature oocytes in the gastrodermal canals, was collected and transported to the laboratory on August 20, 2016 to observe planula development. After planulation, the development of an early planula into a primary polyp was observed under a stereomicroscope and samples for RNA-seq were acquired (Figure 1).

**Figure 1.**
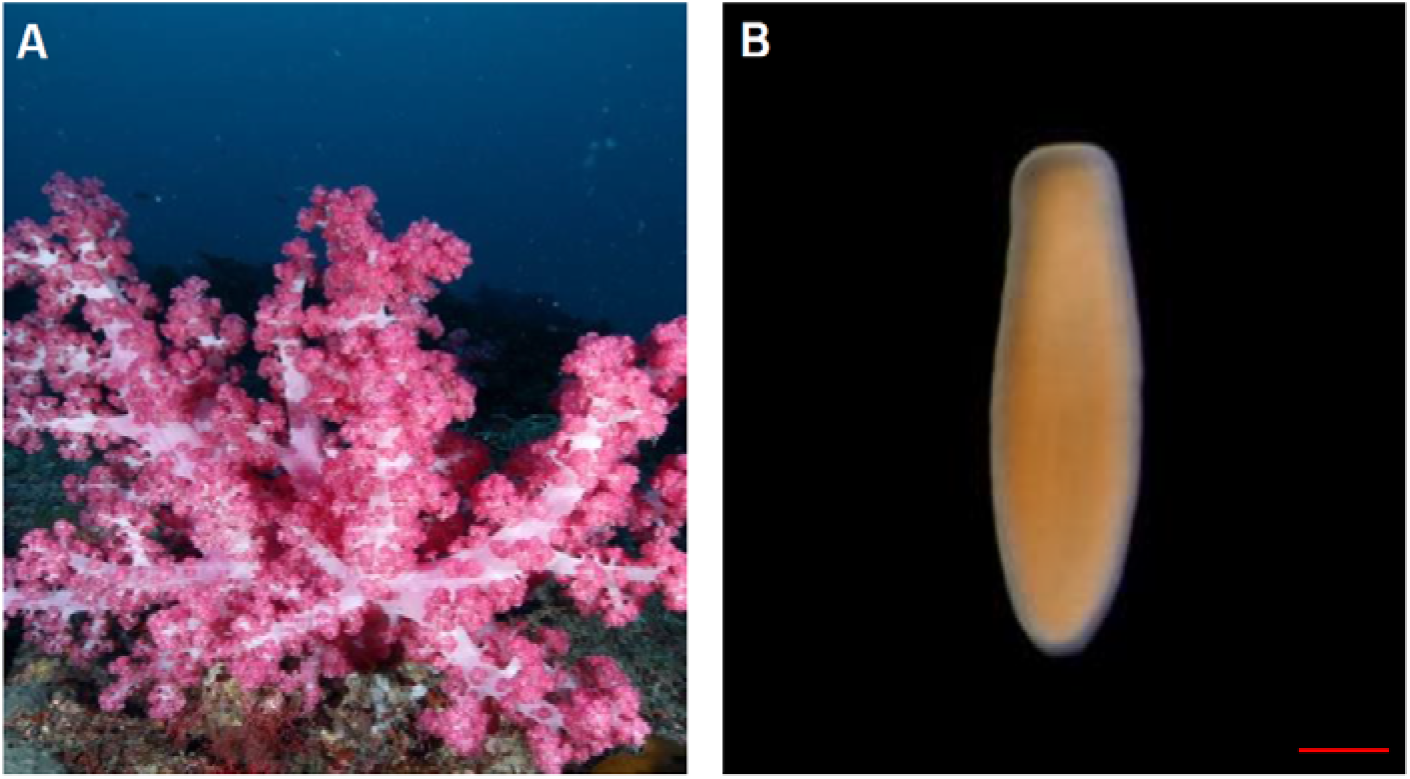
| Adult and larval states of *Dendronephthya gigantea.* **(A)** fully expanded adult colonies. (**B)** a free-swimming planula larva. Red scale bar indicates 200 μm.

For the DNA extraction, the *D. gigantea* colony was mortar-pulverized in liquid nitrogen and the powder homogenized in lysis solution [2% CTAB, 1.4M NaCl, 100 mM Tris-Cl (pH 8.0), 20 mM EDTA, 1% β-mercaptoethanol], and incubated at 65°C for one hour. The same volume of a phenol:chloroform:isoamyl alcohol (23:24:1) mixture was added to denature the proteins and phase separation by centrifugation at 12,000 rpm for 15 min at room temperature was performed. The aqueous phase was retained and incubated at 37°C for one hour after RNase A (30 mg/ml) was added. The DNA was extracted with a phenol:chloroform:isoamyl alcohol (25:24:1) mixture treatment, followed by adding a chloroform:isoamyl alcohol (24:1) mixture with separating phases centrifuged at 10,000g for 15 min at room temperature. In the next step 1/10 volume of 3 M sodium acetate (pH 5.2) and the same volume of 100% ethanol were added into the retained aqueous phase. The precipitated DNA was washed using 70% ethanol and re-suspended in an appropriate volume of ion-exchanged ultrapure water. The DNA quantity was verified by picogreen method using Victor 3 fluorometry and agarose gel electrophoresis.

To extract RNA, the *D. gigantea* whole colony and planula larvae were mortar-pulverized in liquid nitrogen. The tissue powder was then homogenized in 700 μl of lysis solution [35 mM EDTA, 0.7 M LiCl, 7% SDS, 200 mM Tris-Cl (pH 9.0)], and RNA was extracted with 700 μl of water-saturated phenol. A one-third of volume of 8 M LiCl was added into the retained aqueous phase, which was maintained at 4°C for two hours. The RNA was precipitated after centrifugation at 14,000 rpm for 30 min followed by resuspension in 300 μl of DEPC-treated water followed by a reprecipitation with 1/10 volumes of 3 M sodium acetate (pH 5.2) and isopropanol. The precipitated RNA was rinsed with 70% ethanol (diluted in DEPC-treated water) and dissolved in an appropriate volume of DEPC-treated water (30–40 μl). RNA quantity and integrity were analyzed using a NanoDrop ND-1000 spectrometer and an Agilent 2100 Bioanalyzer with RNA Integrity Number (RIN).

### Genome size estimation

We estimated the genome size of *D. gigantea* to be 276 Mb (276,273,039 bp) using 35.54Gb (128-fold coverage) of Illumina short paired-end reads at a k-mer size of 17. The k-mer analysis was conducted using SOAPec (version 2.01) [21]. The graph for the k-mer frequency distribution showed that there were two peaks and the heterozygosity of the *D. gigantea* genome is high [22] (Figure 2). This finding is consistent with previous reports of invertebrates showing relatively high levels of genome heterozygosity [23].

**Figure 2.**
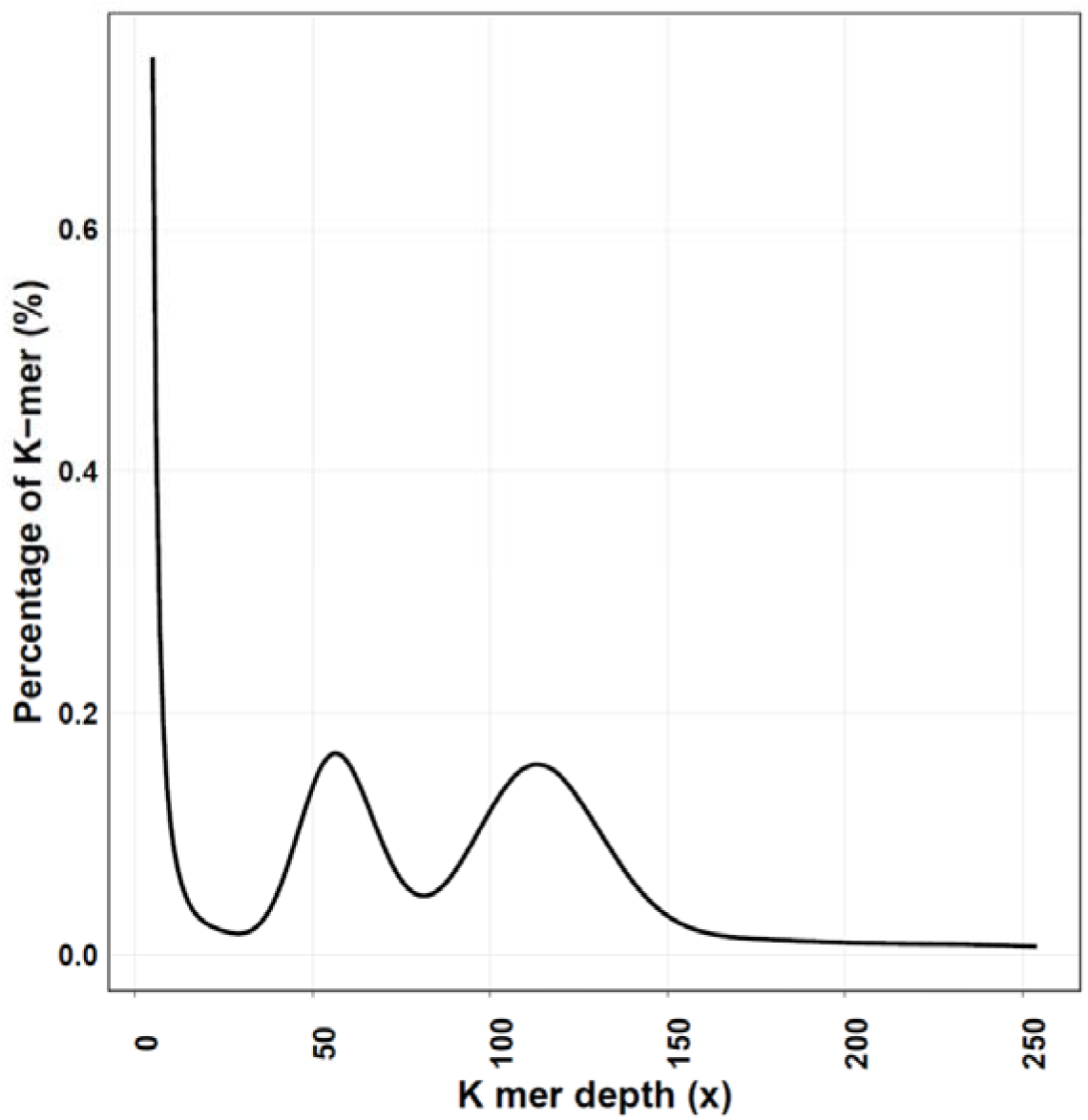
| K-mer (17-mer) frequency percentage distribution curve of sequencing reads of *D. gigantea.* The X-axis represents the k-mer depth (x) and the Y-axis represents the percentage of specific k-mer. There are two peaks in the graph, implying the heterozygosity of the *D. gigantea* genome is high. The left and right peak appear when the k-mer depth is 56 and 113, respectively. The genome size was estimated to be 276 Mb.

### Sequencing and *de novo* genome assembly of the *D. gigantea* genome

We generated 29.85Gb (108-fold coverage) of PacBio long reads for an initial draft assembly which is complemented by 35.54Gb (128-fold coverage) of Illumina short paired-end reads for error-correction. We constructed the first reference genome assembly using *D. gigantea* polyp tissues. It resulted in the longest N50 length (1.4 Mb) reported among cnidarian genomes thus far (Table 1). We filtered out bacterial and fungal DNA reads (about 1.18%) using BLASTN (version 2.2.28) [24] against the UniProt database [25]. The genome was assembled using PacBio long-reads by FALCON (version 0.3.0) [26], with the 10Kb setting in the length cutoff that is used for the seed reads in the initial mapping and pre-assembly. For error-correction, we replaced the assembled contigs of PacBio long-reads with the Illumina short paired-end reads by self-mapping in case of homo-variants and non-reference hetero variants. We repeated this error-correction process three times to correct for sequencing errors to achieve a final contig N50 value of 1,445,523 bp (Table 1).

**Table 1:** Statistics of the *D. gigantea* genome assembly compared to other cnidarians

|  | *Dendronephthya gigantea* | *Orbicella faveolata* | *Stylophora pistillata* | *Acropora digitifera* | *Aiptasia pallida* | *Nematostella vectensis* | *Hydra magnipapillata* |
| --- | --- | --- | --- | --- | --- | --- | --- |
| No. of sequences | 1,323 | 1,933 | 5,688 | 2,421 | 4,312 | 10,804 | 20,916 |
| Total bases (bp) | 286,131,912 | 485,548,939 | 400,120,318 | 447,497,157 | 256,132,296 | 356,613,585 | 852,170,992 |
| Average length (bp) | 216,275 | 251,189 | 70,345 | 184,839 | 59,400 | 33,008 | 40,743 |
| Standard deviation (bp) | 596,503 | 541,789 | 193,436 | 280,650 | 169,768 | 149,438 | 58,784 |
| N50 (bp) | 1,445,523 | 1,162,446 | 457,453 | 483,559 | 442,145 | 472,588 | 96,317 |
| GC contents | 37 % | 39 % | 39 % | 39 % | 36 % | 41 % | 28 % |

### Annotation of repetitive sequences in the *D. gigantea* genome

About 12 % of the *D. gigantea* genome consists of repeat elements. We searched for transposable elements using both *ab initio-* and homology-based methods using RepeatModeler (version 1.0.7) [27] and RepeatMasker (version 4.0.5) [28] and RepeatMasker (version 4.0.5) [28] and Repbase database (version 19.03) [29], respectively. Tandem repeat predictions were performed using Tandem Repeats Finder (version 4.07) [30]. After merging results, we found transposable elements make up an 11.97 % of the *D. gigantea* genome, in which tandem repeats and long terminal repeat elements (LTR) represented 7.24% and 2.25% of the genome, respectively (Table 2).

**Table 2:** Repeat sequences in the *D. gigantea* genome.

| Repeat type | Ab initio based (bp) | Homology based (bp) | Total (bp) | Percentage of genome (%) |
| --- | --- | --- | --- | --- |
| DNA | 5,989,055 | 2,444,506 | 6,344,179 | 2.22 |
| LINE | 2,621,991 | 1,893,795 | 3,014,162 | 1.05 |
| LTR | 6,186,765 | 4,707,866 | 6,435,444 | 2.25 |
| Low complexity | 36,863 | 41,827 | 42,373 | 0.015 |
| SINE | 4,753 | - | 4,753 | 0.0017 |
| Satellite | 244,013 | 9,371 | 244,167 | 0.085 |
| Simple repeat | 619,655 | 1,721,644 | 1,727,993 | 0.60 |
| Tandem repeat* | - | - | 20,729,359 | 7.24 |
| Unknown | 142,961 | 242,937 | 253,480 | 0.09 |
| Unspecified | 2,153,035 | - | 2,153,035 | 0.75 |
| Total transposable elements | 16,760,059 | 10,828,627 | 34,254,188 | 11.97<sup>§</sup> |

\* Tandem repeats were separately predicted using Tandem Repeats Finder (version 4.07) [30].
<sup>§</sup> The total element sum is smaller than the arithmetic sum of the repeat types because there

### Gene prediction, annotation, and quality assessment

We found just under 29,000 protein-coding genes in *D. gigantea* (Table 3). We selected our final gene set after comparing two methods. First, we merged *ab initio-* and homology-based predictions using AUGUSTUS (version 3.1) [31–37] with additional information obtained from homology-based predicted *D. gigantea* gene models, RNA-seq data of the planula and polyp of *D. gigantea* and polyps of *Scleronephthya gracillimum* (unpublished data), and Expressed Sequence Tags (ESTs) of corals downloaded from NCBI database [38]. We used homology-based methods to align repeat-masked *D. gigantea* genome to proteomes of cnidarians obtained from the UniProt database [25], *H. sapiens, M. musculus,* and *D. rerio* using GeneBlastA (version 1.0.4) [39] with E-value cutoff 1E-05 and Exonerate (version 2.2.0) [40]. We gained 8,669 gene models from the homology-based method and these were used as exon hints when we merged both of the *ab initio-* and homology-based methods. In addition, we aligned RNA-seq reads to the *D. gigantea* genome using TopHat (version 2.0.9) [41] to use as intron hints. The EST sequences of corals were mapped to the genome assembly using BLAT (version 34) [42] and used as exon and intron hints. The gene models were filtered according to these criteria: final gene models must both start and stop codons, CDS length is a multiple of three, and the length of protein-coding genes is more than 40 amino acids. In addition, single exon genes with FPKM value < 1 were filtered out when multiple exons existed with the same gene symbol from the UniProt database [25]. After filtering, 28,879 protein-coding gene predictions remained (Table 3).

**Table 3:** Statistics of protein-coding genes in *D. gigantea*

|  | Number | Percentage (%) |
| --- | --- | --- |
| Pre-filtered gene models | 32,487 | 100.00 |
| Gene models with amino acid length > 40 | 32,478 | 99.97 |
| Gene models whose CDS length is multiple of 3 | 32,256 | 99.29 |
| Complete gene models containing both start and stop codons | 32,150 | 98.96 |
| Single exon genes with FPKM value < 1 when multi exon gene exist as same symbol | 3,310 | 10.19 |
| Total number of final gene models | 28,879 | 88.89 |

In a second approach, we combined predicted genes from the Maker pipeline (version 2.31.10) [43] with those from BRAKER2 (version 2.1.2) [31, 33, 44]. To obtain additional evidence for predicted genes, we mapped assembled transcripts from RNA-seq data of the planula and polyp of *D. gigantea,* sequences from Swiss-Prot database [25], and genes from closely related species to the *D. gigantea* genome using BLAST (version 2.2.28) [24] and Exonerate (version 2.2.0) [40]. The predicted genes with less than 1.00 AED score were sorted as the final set. This gave 28,937 protein-coding genes as best gene models by comparing the genes from the BRAKER2 [31, 33, 44] with those from the Maker pipeline [43].

We compared both gene sets using BUSCO [45, 46] which provides quality estimation by the number of predicted orthologs, which showed comparable high quality, increasing our confidence in our set of predicted genes. Afterwards, we finalized the gene set obtained by the first method because the BUSCO (version 3.0.2) [45, 46] assessments showed that gene set from the first method (93.97% complete BUSCO genes) had a slightly better quality than that of the second method (93.35% complete BUSCO genes) (Table 4).

**Table 4:** Comparison of BUSCO assessments of the gene sets between two methods

|  | Current gene set |  | Not current gene set |  |
| --- | --- | --- | --- | --- |
|  | First method |  | Second method |  |
|  | Number | Percentage (%) | Number | Percentage (%) |
| Complete single copy BUSCO genes | 854 | 87.32 | 806 | 82.41 |
| Complete duplicated BUSCO genes | 65 | 6.65 | 107 | 10.94 |
| Complete BUSCO genes (single copy + duplicated) | 919 | 93.97 | 913 | 93.35 |
| Fragmented BUSCO genes | 24 | 2.45 | 35 | 3.58 |
| Missing BUSCO genes | 35 | 3.58 | 30 | 3.07 |
| Total of used genes in BUSCO | 978 | - | 978 | - |

*D. gigantea* had high gene set quality among the cnidarians covering about 94% of the complete BUSCO ortholog benchmark genes (Figure 3). We compared the quality of the *D. gigantea* gene models with six published cnidarians *(Aiptasia pallida, Acropora digitifera, Hydra magnipapillata, Nematostella vectensis, Orbicella faveolata,* and *Stylophora pistillata)* using BUSCO (version 3.0.2) [45, 46]. The *D. gigantea* gene models had the highest number of the complete single copy BUSCO genes; 87.32% complete single copy BUSCO genes among cnidarians (Figure 3). It also had the second highest value of complete BUSCO genes which included both single copy and duplicated genes among cnidarians. (Figure 3).

**Figure 3.**
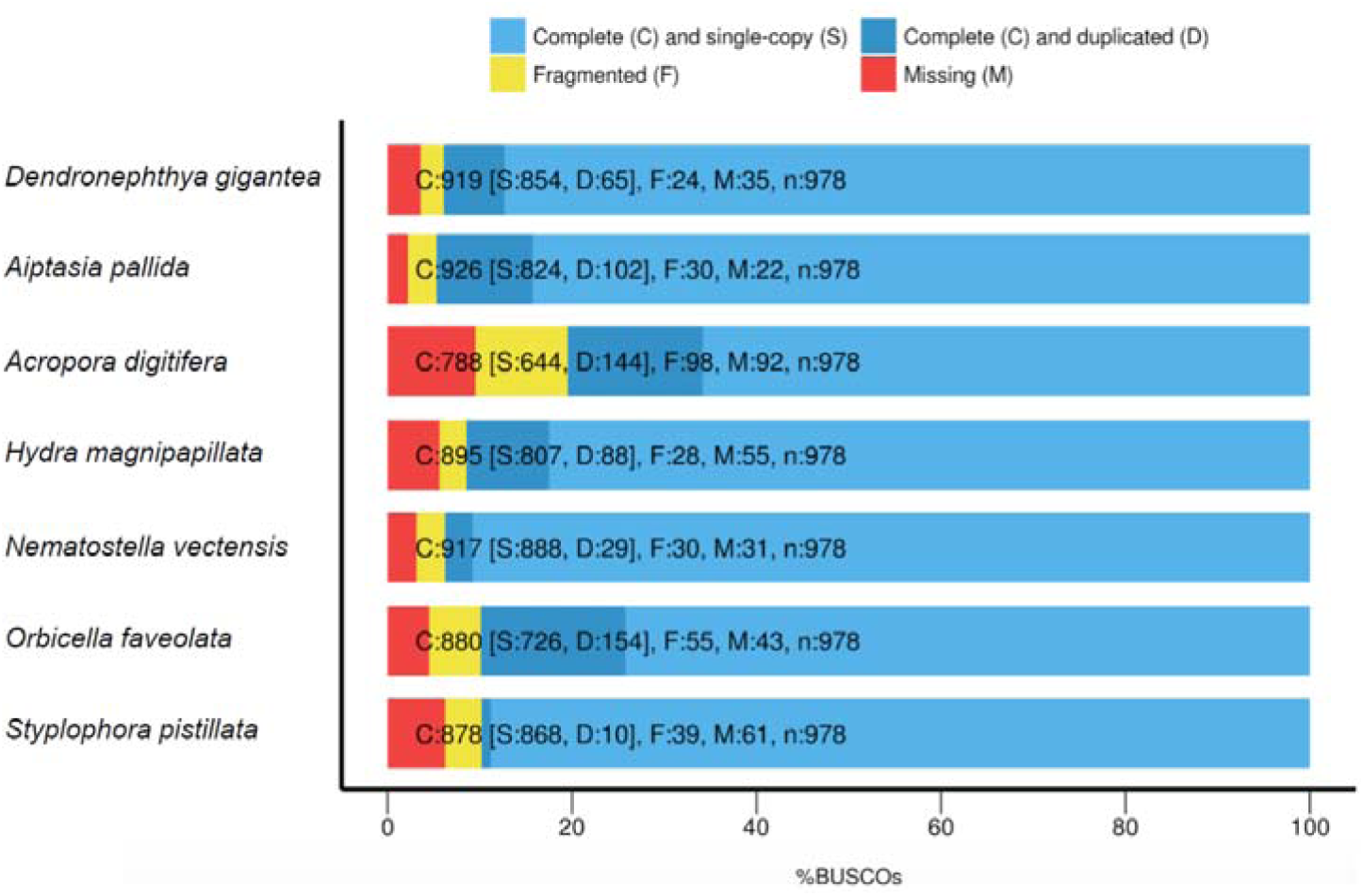
| Assessment of the *D. gigantea* gene models compared to other cnidarians. The figure shows results of BUSCO analysis. Light-blue denotes the complete single-copy genes, dark-blue denotes complete duplicated genes, yellow denotes fragmented genes, and red denotes missing genes.

### Phylogenetic analysis and species divergence time estimation

We found that *D. gigantea* has diverged the earliest among the anthozoans based on our calculation as follows. First, we examined orthologous gene clustering of complete protein-coding genes from the six published cnidarians *(Orbicella faveolata, Stylophora pistillata, Acropora digitifera, Nematostella vectensis, Aiptasia pallida,* and *Hydra magnipapillata)* and seven non-cnidarian metazoans (*Danio rerio*, *Homo Sapiens*, *Drosophila melanogaster*, *Caenorhabditis elegans, Trichoplax adhaerens, Amphimedon queenslandica,* and *Mnemiopsis leidyi).* Our out-group was the unicellular holozoan, *Monosiga brevicollis.* Clusters were generated using OrthoMCL (version 2.0.9) [47] with an E-value cutoff of 1E-20. We found that *D. gigantea* contains 12,597 orthologous gene families, excluding singletons, of which 3,656 are shared with stony corals *(Orbicella faveolata, Stylophora pistillata,* and *Acropora digitifera)* and hydra *(Hydra magnipapillata)* (Figure 4). A total of 4,863 gene families were specific to the *D. gigantea* (Figure 4). Secondly, molecular phylogenetic analysis suggested the divergence of the octocoral *(D. gigantea)* and other three stony corals *(O. faveolata, S. pistillata,* and *A. digitifera)* happened 544 million years ago (Figure 5). Finally, we estimated the phylogeny by using 197 single copy orthologs using PROTGAMMAJTT model in RAxML (version 8.2.8) [48]. The divergence times were estimated by MCMCtree program in PAML package (version 4.8) [49] with the independent rates model (clock=2). The date of the node between *D. melanogaster-C. elegans* was constrained to 743 MYA and *H. sapiens-D. rerio* was constrained to 435 MYA based on the TimeTree database [50]. Notably, our results show that the octocoral, *D. gigantea*, is located between hexacorallia and hydrozoa (Figure 5), implying that the octocoral is the earliest diverged group among anthozoans.

**Figure 4.**
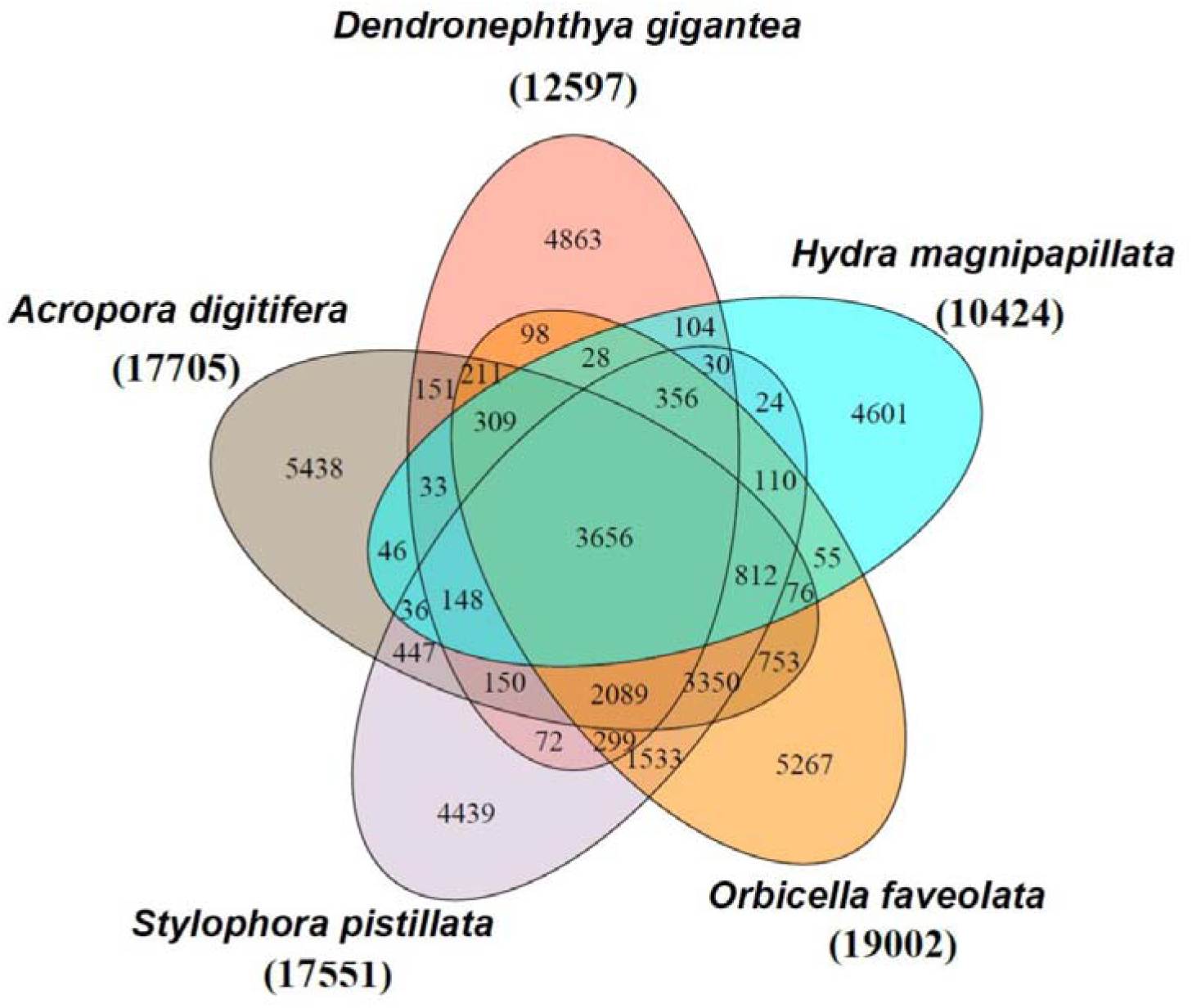
| A Venn-diagram of orthologous gene families. The Venn-diagram shows shared and specific gene families in the *D. gigantea, A. digitifera, S. pistillata, O. faveolata,* and *H. magnipapillata* genomes. The total numbers of gene families are given in parentheses.

**Figure 5.**
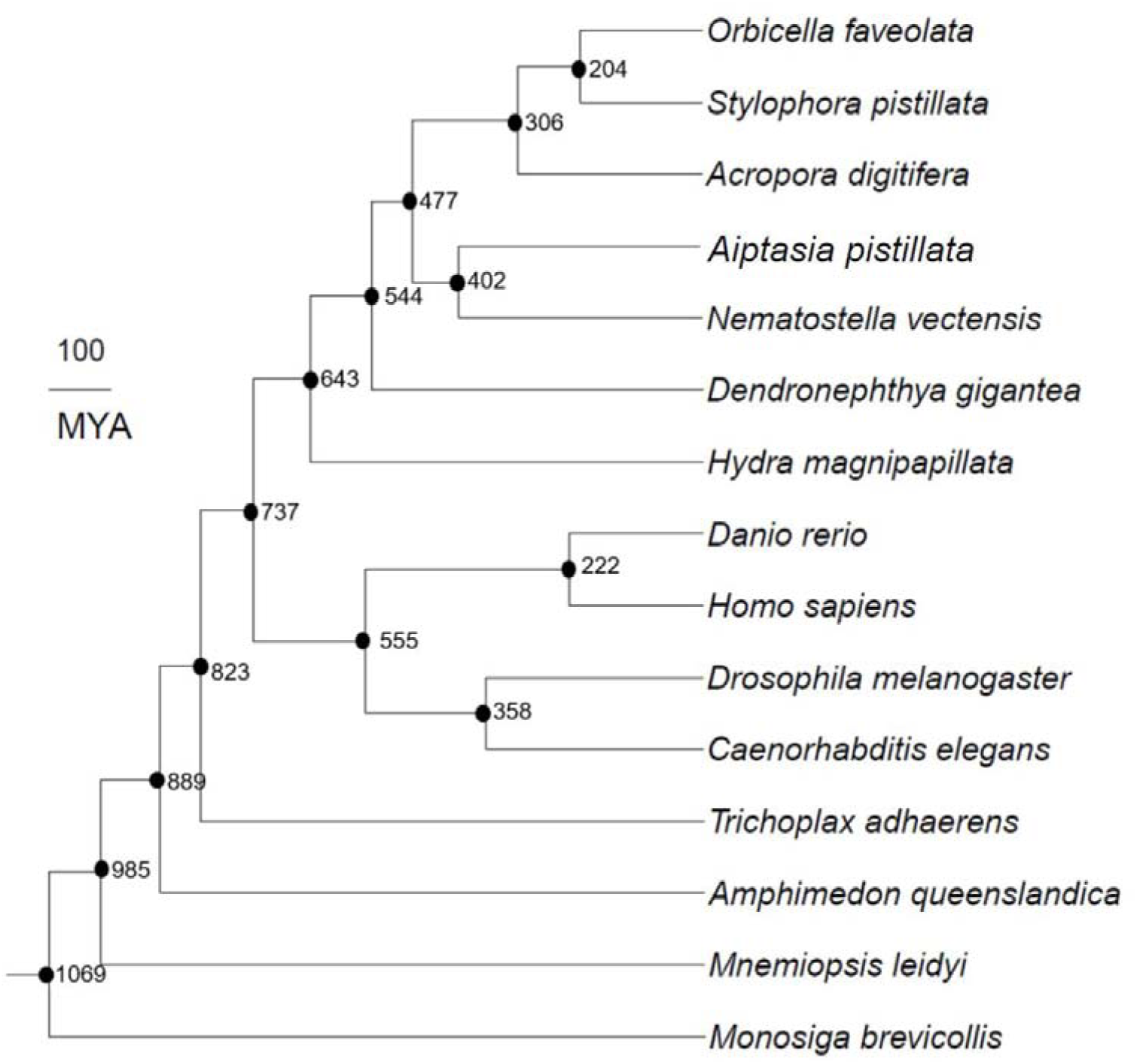
| Phylogenetic relationship of *D. gigantea* with other species. Tree shows the phylogeny with divergence time among 15 species. Numbers in each branch denote the estimated divergence time (million years ago).

### Hox gene clusters in cnidarians

Analyses of Hox (homeobox) genes revealed differences between the soft and stony corals. Hox genes encode transcription factors that perform diverse roles during development [51]. They are best known to define body plan [51]. To identify and classify Hox gene clusters, we found all instances of the homeobox domain based on Pfam database [52] using HMMER (version 3.1b2) [53] and InterProScan (version 5.32–71.0) [54, 55]. Genes with the homeobox domain were classified using BLAST (version 2.2.28) [24] against HomeoDB [56, 57] and mapping to the homeobox domain of *N. vectensis* Hox genes from GenBank [38]. We found the three stony corals have a similar and familiar pattern of Hox gene clusters [18] (Figure 6). However, Evx which is a member of the Antp superclass of Hox genes [58] is absent in *D. gigantea* (Figure 6) which needs to be verified by experiments.

**Figure 6.**
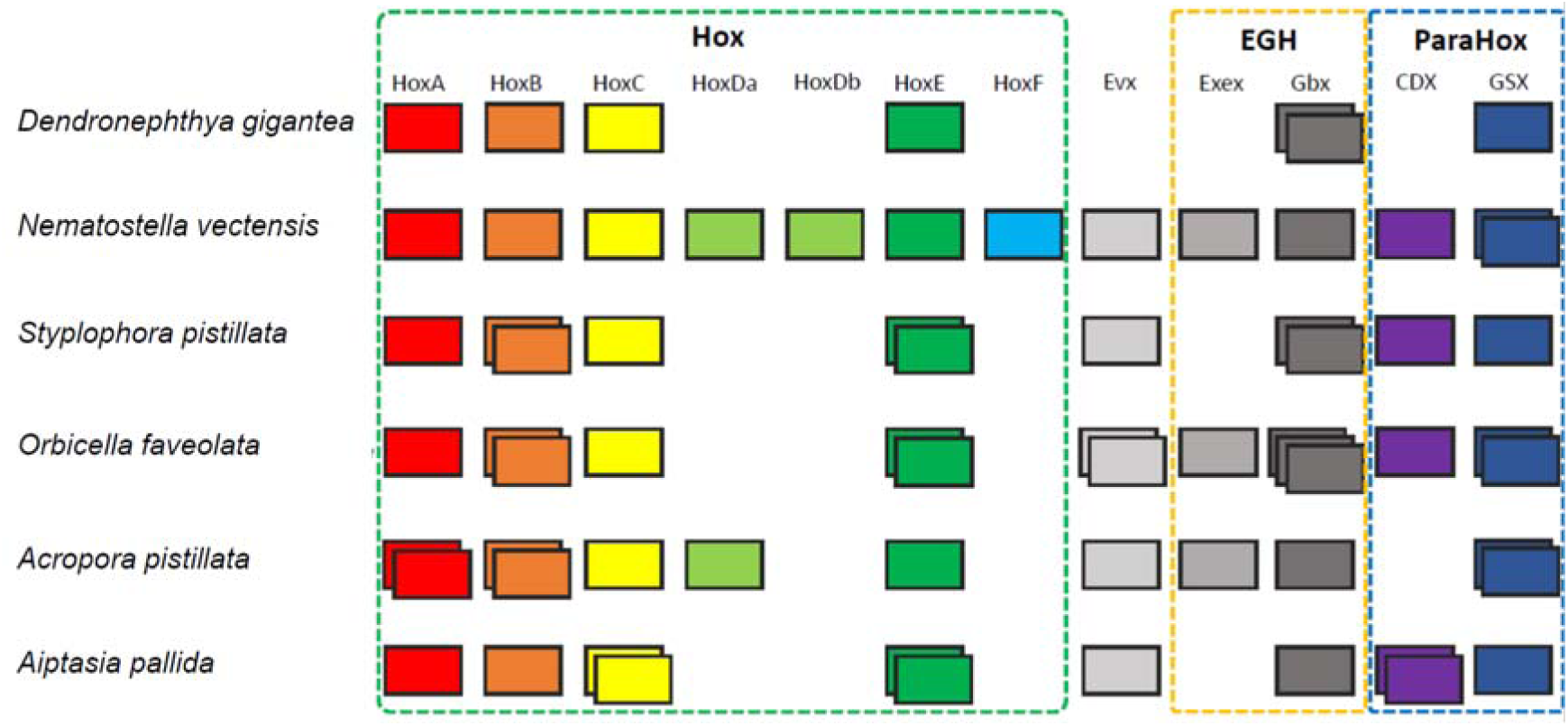
| Hox gene clusters of *D. gigantea* and other anthozoans. Green dashed-line box denotes Hox gene cluster (HoxA, HoxB, HoxC, HoxDa, HoxDb, HoxE, and HoxF), yellow dashed-line box denotes EGF gene cluster (Evex and Gbx), and blue dashed-line box denotes ParaHox gene cluster (CDX and GSX). The number of boxes shows the number of each gene copies in the genome.

### Conclusion

We present a high-quality, draft genome from the non-symbiotic octocoral, *Dendronephthya gigantea* with which we find that the octocoral is the earliest diverged group among anthozoans showing the divergence time estimation of 544 million years from the stony corals. It adds a new octocoral assembly for cnidarians, in addition to hexacoral and hydra genomes, thus opening the doors to in depth comparative analyses of stony and soft corals and symbiotic and non-symbiotic coral genomes. Furthermore, future study of the genome and transcriptome set we provide here may contribute new answers about the relative successes of genetic coping mechanisms between soft and stony corals in terms of calcification and survival strategies in the face of global warming and ocean acidification.

### Availability of Data and Materials

Raw DNA and RNA sequencing data and genome assembly are available at NCBI under the project accession number PRJNA507923 and PRJNA507943.

## Abbreviations

MYA: million years ago
EST: Expressed Sequence Tag
BUSCO: Benchmarking Universal Single-Copy Orthologs
FPKM: Fragments Per Kilobase Million
AED: Annotation Edit Distance

## Acknowledgements

Thanks to the Korea Institute of Science and Technology Information (KISTI) for providing us access to the Korea Research Environment Open NETwork (KREONET), an internet connection service supporting efficient information and data transfer.

## Author’s contributions

J.B. and S.Y. supervised the project. Y.S.C., J.B., and S.Y. planned and coordinated the project. Yeonsu J., J.A.W., A.B., S.Y. and J.B. wrote the manuscript. Nayun L., S.J.H., Nayoung L., Yejin J., S.W., J.H.L., H.S.Y and S.Y. prepared the samples, performed the experiments. Yeonsu J., S.G.P, H.S.K., H.M.K., Y.B., S.J., T.R., H.K., and Y.S.C. performed in-depth bioinformatics data analyses. All authors reviewed the manuscript and discussed the work.

## References

1. Friedlander AM, Parrish JD: Habitat characteristics affecting fish assemblages on a Hawaiian coral reef. Journal of Experimental Marine Biology and Ecology 1998, 224: 1–30.

2. van de Water JA, Allemand D, Ferrier-Pagès C: Host-microbe interactions in octocoral holobionts-recent advances and perspectives. Microbiome 2018, 6: 64.

3. Gili J-M, Coma R: Benthic suspension feeders: their paramount role in littoral marine food webs. Trends in ecology & evolution 1998, 13: 316–321.

4. Carpenter KE, Abrar M, Aeby G, Aronson RB, Banks S, Bruckner A, Chiriboga A, Cortés J, Delbeek JC, DeVantier L: One-third of reef-building corals face elevated extinction risk from climate change and local impacts. Science 2008.

5. Hoegh-Guldberg O: Climate change, coral bleaching and the future of the world’s coral reefs. Marine and freshwater research 1999, 50: 839–866.

6. de Paula AF, Creed JC: Two species of the coral Tubastraea (Cnidaria, Scleractinia) in Brazil: a case of accidental introduction. Bulletin of Marine Science 2004, 74: 175–183.

7. Santodomingo N, Reyes J, Florez P, Chacón-Gómez IC, van Ofwegen LP, Hoeksema BW: Diversity and distribution of azooxanthellate corals in the Colombian Caribbean. Marine Biodiversity 2013, 43: 7–22.

8. Inoue S, Kayanne H, Yamamoto S, Kurihara H: Spatial community shift from hard to soft corals in acidified water. Nature Climate Change 2013, 3: 683.

9. Pandolfi JM, Connolly SR, Marshall DJ, Cohen AL: Projecting coral reef futures under global warming and ocean acidification. science 2011, 333: 418–422.

10. Ries J, Cohen A, McCorkle D: A nonlinear calcification response to CO 2-induced ocean acidification by the coral Oculina arbuscula. Coral Reefs 2010, 29: 661–674.

11. de Putron SJ, McCorkle DC, Cohen AL, Dillon A: The impact of seawater saturation state and bicarbonate ion concentration on calcification by new recruits of two Atlantic corals. Coral Reefs 2011, 30: 321–328.

12. Lopes AR, Faleiro F, Rosa IC, Pimentel MS, Trubenbach K, Repolho T, Diniz M, Rosa R: Physiological resilience of a temperate soft coral to ocean warming and acidification. Cell Stress and Chaperones 2018: 1–8.

13. Shinzato C, Shoguchi E, Kawashima T, Hamada M, Hisata K, Tanaka M, Fujie M, Fujiwara M, Koyanagi R, Ikuta T: Using the Acropora digitifera genome to understand coral responses to environmental change. Nature 2011, 476: 320.

14. Baumgarten S, Simakov O, Esherick LY, Liew YJ, Lehnert EM, Michell CT, Li Y, Hambleton EA, Guse A, Oates ME: The genome of Aiptasia, a sea anemone model for coral symbiosis. Proceedings of the National Academy of Sciences 2015, 112: 11893–11898.

15. Putnam NH, Srivastava M, Hellsten U, Dirks B, Chapman J, Salamov A, Terry A, Shapiro H, Lindquist E, Kapitonov VV: Sea anemone genome reveals ancestral eumetazoan gene repertoire and genomic organization. science 2007, 317: 86–94.

16. Voolstra CR, Li Y, Liew YJ, Baumgarten S, Zoccola D, Flot J-F, Tambutté S, Allemand D, Aranda M: Comparative analysis of the genomes of Stylophora pistillata and Acropora digitifera provides evidence for extensive differences between species of corals. Scientific reports 2017, 7: 17583.

17. Snelling J, Dziedzic K, Guermond S, Meyer E: Development of an integrated genomic map for a threatened Caribbean coral (Orbicella faveolata). 2017.

18. Ying H, Cooke I, Sprungala S, Wang W, Hayward DC, Tang Y, Huttley G, Ball EE, Forêt S, Miller DJ: Comparative genomics reveals the distinct evolutionary trajectories of the robust and complex coral lineages. Genome biology 2018, 19: 175.

19. Hwang S-J, Song J-I: Reproductive biology and larval development of the temperate soft coral Dendronephthya gigantea (Alcyonacea: Nephtheidae). Marine Biology 2007, 152: 273–284.

20. Imbs AB, Latyshev NA, Zhukova NV, Dautova TN: Comparison of fatty acid compositions of azooxanthellate Dendronephthya and zooxanthellate soft coral species. Comparative Biochemistry and Physiology Part B: Biochemistry and Molecular Biology 2007, 148: 314–321.

21. Luo R, Liu B, Xie Y, Li Z, Huang W, Yuan J, He G, Chen Y, Pan Q, Liu Y: SOAPdenovo2: an empirically improved memory-efficient short-read de novo assembler. Gigascience 2012, 1: 18.

22. Liu B, Shi Y, Yuan J, Hu X, Zhang H, Li N, Li Z, Chen Y, Mu D, Fan W: Estimation of genomic characteristics by analyzing k-mer frequency in de novo genome projects. arXiv preprint arXiv:13082012 2013.

23. Ellegren H, Galtier N: Determinants of genetic diversity. Nature Reviews Genetics 2016, 17: 422.

24. Altschul SF, Gish W, Miller W, Myers EW, Lipman DJ: Basic local alignment search tool. Journal of molecular biology 1990, 215: 403–410.

25. Consortium U: UniProt: the universal protein knowledgebase. Nucleic acids research 2018, 46: 2699.

26. Chin C-S, Peluso P, Sedlazeck FJ, Nattestad M, Concepcion GT, Clum A, Dunn C, O'Malley R, Figueroa-Balderas R, Morales-Cruz A: Phased diploid genome assembly with single-molecule real-time sequencing. Nature methods 2016, 13: 1050.

27. Price AL, Jones NC, Pevzner PA: De novo identification of repeat families in large genomes. Bioinformatics 2005, 21:i351-i358.

28. Chen N: Using RepeatMasker to identify repetitive elements in genomic sequences. Current protocols in bioinformatics 2004, 5:4.10. 11–14.10. 14.

29. Jurka J, Kapitonov VV, Pavlicek A, Klonowski P, Kohany O, Walichiewicz J: Repbase Update, a database of eukaryotic repetitive elements. Cytogenetic and genome research 2005, 110: 462–467.

30. Benson G: Tandem repeats finder: a program to analyze DNA sequences. Nucleic acids research 1999, 27: 573–580.

31. Stanke M, Diekhans M, Baertsch R, Haussler D: Using native and syntenically mapped cDNA alignments to improve de novo gene finding. Bioinformatics 2008, 24: 637–644.

32. Stanke M, Tzvetkova A, Morgenstern B: AUGUSTUS at EGASP: using EST, protein and genomic alignments for improved gene prediction in the human genome. Genome biology 2006, 7:S11.

33. Stanke M, Schöffmann O, Morgenstern B, Waack S: Gene prediction in eukaryotes with a generalized hidden Markov model that uses hints from external sources. BMC bioinformatics 2006, 7: 62.

34. Stanke M, Morgenstern B: AUGUSTUS: a web server for gene prediction in eukaryotes that allows user-defined constraints. Nucleic acids research 2005, 33:W465-W467.

35. Stanke M, Steinkamp R, Waack S, Morgenstern B: AUGUSTUS: a web server for gene finding in eukaryotes. Nucleic acids research 2004, 32: W309–W312.

36. Stanke M, Waack S: Gene prediction with a hidden Markov model and a new intron submodel. Bioinformatics 2003, 19:ii215-ii225.

37. Stanke M: Gene prediction with a hidden Markov model. 2004.

38. Benson DA, Cavanaugh M, Clark K, Karsch-Mizrachi I, Ostell J, Pruitt KD, Sayers EW: GenBank. Nucleic acids research 2017.

39. She R, Chu JS-C, Wang K, Pei J, Chen N: GenBlastA: enabling BLAST to identify homologous gene sequences. Genome research 2009, 19: 143–149.

40. Slater GSC, Birney E: Automated generation of heuristics for biological sequence comparison. BMC bioinformatics 2005, 6: 31.

41. Trapnell C, Pachter L, Salzberg SL: TopHat: discovering splice junctions with RNA-Seq. Bioinformatics 2009, 25: 1105–1111.

42. Kent WJ: BLAT—the BLAST-like alignment tool. Genome research 2002, 12: 656–664.

43. Cantarel BL, Korf I, Robb SM, Parra G, Ross E, Moore B, Holt C, Alvarado AS, Yandell M: MAKER: an easy-to-use annotation pipeline designed for emerging model organism genomes. Genome research 2008, 18: 188–196.

44. Hoff KJ, Lange S, Lomsadze A, Borodovsky M, Stanke M: BRAKER1: unsupervised RNA-Seq-based genome annotation with GeneMark-ET and AUGUSTUS. Bioinformatics 2015, 32: 767–769.

45. Waterhouse RM, Seppey M, Simão FA, Manni M, Ioannidis P, Klioutchnikov G, Kriventseva EV, Zdobnov EM: BUSCO applications from quality assessments to gene prediction and phylogenomics. Molecular biology and evolution 2017, 35: 543–548.

46. Simão FA, Waterhouse RM, Ioannidis P, Kriventseva EV, Zdobnov EM: BUSCO: assessing genome assembly and annotation completeness with single-copy orthologs. Bioinformatics 2015, 31: 3210–3212.

47. Li L, Stoeckert CJ, Roos DS: OrthoMCL: identification of ortholog groups for eukaryotic genomes. Genome research 2003, 13: 2178–2189.

48. Stamatakis A: RAxML version 8: a tool for phylogenetic analysis and post-analysis of large phylogenies. Bioinformatics 2014, 30: 1312–1313.

49. Yang Z: PAML 4: phylogenetic analysis by maximum likelihood. Molecular biology and evolution 2007, 24: 1586–1591.

50. Kumar S, Stecher G, Suleski M, Hedges SB: TimeTree: a resource for timelines, timetrees, and divergence times. Molecular Biology and Evolution 2017, 34: 1812–1819.

51. Akam M: Hox genes and the evolution of diverse body plans. Phil Trans R Soc Lond B 1995, 349: 313–319.

52. Finn RD, Coggill P, Eberhardt RY, Eddy SR, Mistry J, Mitchell AL, Potter SC, Punta M, Qureshi M, Sangrador-Vegas A: The Pfam protein families database: towards a more sustainable future. Nucleic acids research 2015, 44:D279-D285.

53. Finn RD, Clements J, Eddy SR: HMMER web server: interactive sequence similarity searching. Nucleic acids research 2011, 39: W29–W37.

54. Mitchell AL, Attwood TK, Babbitt PC, Blum M, Bork P, Bridge A, Brown SD, Chang H-Y, El-Gebali S, Fraser MI: InterPro in 2019: improving coverage, classification and access to protein sequence annotations. Nucleic acids research 2018.

55. Jones P, Binns D, Chang H-Y, Fraser M, Li W, McAnulla C, McWilliam H, Maslen J, Mitchell A, Nuka G: InterProScan 5: genome-scale protein function classification. Bioinformatics 2014, 30: 1236–1240.

56. Zhong Yf, Holland PW: HomeoDB2: functional expansion of a comparative homeobox gene database for evolutionary developmental biology. Evolution & development 2011, 13: 567–568.

57. Zhong YF, Butts T, Holland PW: HomeoDB: a database of homeobox gene diversity. Evolution & development 2008, 10: 516–518.

58. Patel NH, Prince VE: Beyond the Hox complex. Genome Biology 2000, 1:reviews1027. 1021.

